# Sexual Differences in Mitochondrial Proteins in Rat Cerebral Microvessels: A Proteomic Approach

**DOI:** 10.1101/2019.12.17.879254

**Authors:** Sinisa Cikic, Partha K. Chandra, Jarrod C. Harman, Ibolya Rutkai, Prasad V.G. Katakam, Jessie J. Guidry, Jeffrey M. Gidday, David W. Busija

## Abstract

Sex differences in mitochondrial numbers and function are present in large cerebral arteries, but it is unclear whether these differences extend to the microcirculation. We performed an assessment of mitochondria-related proteins in cerebral microvessels (MVs) isolated from young, male and female, Sprague-Dawley rats. MVs composed of arterioles, capillaries, and venules were isolated from the cerebrum and used to perform a 3 vs. 3 quantitative, multiplexed proteomics experiment utilizing tandem mass tags (TMT), coupled with liquid chromatography/mass spectrometry (LC/MS). MS data and bioinformatic analyses were performed using Proteome Discoverer version 2.2 and Ingenuity Pathway Analysis. We identified a total of 1,969 proteins, of which 1,871 were quantified by TMT labels. Sixty-four proteins were expressed significantly (p < 0.05) higher in female samples compared with male samples. Females expressed more mitochondrial proteins involved in energy production, mitochondrial membrane structure, anti-oxidant enzyme proteins, and those involved in fatty acid oxidation. Conversely, males had higher expression levels of mitochondria-destructive proteins. We validated our key Proteomics results with western blotting. Our findings reveal, for the first time, the full extent of sexual dimorphism in the mitochondrial metabolic protein profiles of MVs, which may contribute to sex-dependent cerebrovascular and neurological pathologies.

**Synopsis:** Energy-producing proteins in the cerebral microvessels (MVs) of male and female rats were examined by quantitative discovery-based proteomics to gain insight into the sex-dependent etiology of cardiovascular and neurological diseases. Females expressed more mitochondrial proteins involved in energy production, membrane structure, anti-oxidant activity, and fatty acid oxidation. In contrast, males exhibited more mitochondria-destructive proteins such as mitochondrial eating protein. Our findings reveal for the first time the sexual dimorphism of mitochondria-related proteins in cerebral MVs, which may explain functional sex-related differences in MVs during health and in the etiology of neurological pathologies of cerebrovascular origin.

## INTRODUCTION

Mitochondria generate ATP via oxidative phosphorylation (OXPHOS) and produce other biologically important substances including mitochondrial reactive oxygen species (mitoROS) and small peptides (mitopeptides) (Busija et al, 2016; Yen et al, 2013). Mitochondria are dynamic and change numbers (mitogenesis/mitophagy), shape (fission/fusion), location, and rapidly alter respiration, mitoROS release, anti-oxidant capacity, and mitopeptide production in relation to physiological status (Sure et al, 2018; Bachar et al, 2010; Busija & Katakam 2014; Busija et al, 2016; Dai et al, 2012; Ungvari et al, 2010; Reddy & Beal 2008; Grimmig et al, 2017; Yen et al, 2013; Yu et al, 2017). Previous studies have shown that mitochondrial numbers and DNA, mitochondrial mediated signaling, and mitochondrial respiration is greater in females when compared to males large cerebral arteries in rats (Rutkai et al, 2015; de Souza Mota et al, 2017; Díaz et al, 2019). Also, mitochondrial responses to pathological events such as experimental strokes differ in the large cerebral arteries of male and female rats (Rutkai et al, 2017, Demarest et al, 2016; Netto et al. 2017; Chauhan et al, 2017). However, it is unclear whether sex-dependent differences extend to the cerebral microcirculation. Cerebral microvessels (MVs) are more susceptible to injury than large arteries during aging, injury, and pathology and recent evidence implicates derangement of the microcirculation as a major contributor to the development of neurological diseases such as cognitive impairment (Lee et al, 2019a; Iadecola et al, 2019), vascular dementia (van Leijsen et al, 2017; Wardlaw et al, 2019), and Alzheimer’s disease (AD) (Hansra et al, 2019). Although the mechanisms linking cerebral MV disease with neurological diseases are not fully known, mitochondria in the microcirculation appear to play a central role in disease progression and present an accessible, possible target for sex-specific therapies.

Traditional methodologies, such as Western blotting, restrict the number of mitochondrial-related proteins in cerebral blood vessels that could be efficiently studied at one time and require a *priori* decisions concerning protein selection. However, newer technologies such as proteomics allow large-scale analyses of proteins, including relative abundance and functional inter-relationships, on the same samples (Brzica et al, 2018; Harman et al, 2018; Javitt & Marbl 2019; Tharakan et al, 2019;Vandenbrouck et al, 2019). Although proteomics approaches have been applied to the field of neuroscience (Gomez-Zepeda et al, 2019; Porte et al, 2017), to date there are no reported studies of sex-dependent differences in mitochondria-related proteins in brain MVs.

Therefore, we conducted a comprehensive proteomic characterization of sex-dependent differences in the brain MVs of Sprague-Dawley rats, focusing specifically on mitochondria-related proteins. Using a quantitative discovery-based mass spectrometry (MS) approach, we identified and quantified 1,871 proteins, of which 64 were expressed significantly more in female samples. We validated our key Proteomics results with western blotting. In addition, our findings provide a foundation for future research in fundamental brain MV physiology and pathophysiology (Porte et al, 2017), and contribute to the tissue-specific database necessary to advance algorithm-based bioinformatic platforms such as Ingenuity Pathway Analysis (IPA), that give context and meaning to “big data”/omic experiments.

## RESULTS

### MV composition

Microscopic examination of isolated MVs from male and female rats revealed no gross contamination with non-vascular tissue elements. High magnification (20X) light microscopy revealed that the MV preparation contained a mixture of end arterioles, capillaries, and venules with MV similar preparation characteristics in both males and females (Fig. 1A). Both unstained and alkaline phosphatase (AP)-stained MV segments were detected (Fig. 1B), confirming that the isolated MVs were a mixture of arterioles, capillaries, and venules.

**Fig. 1:**
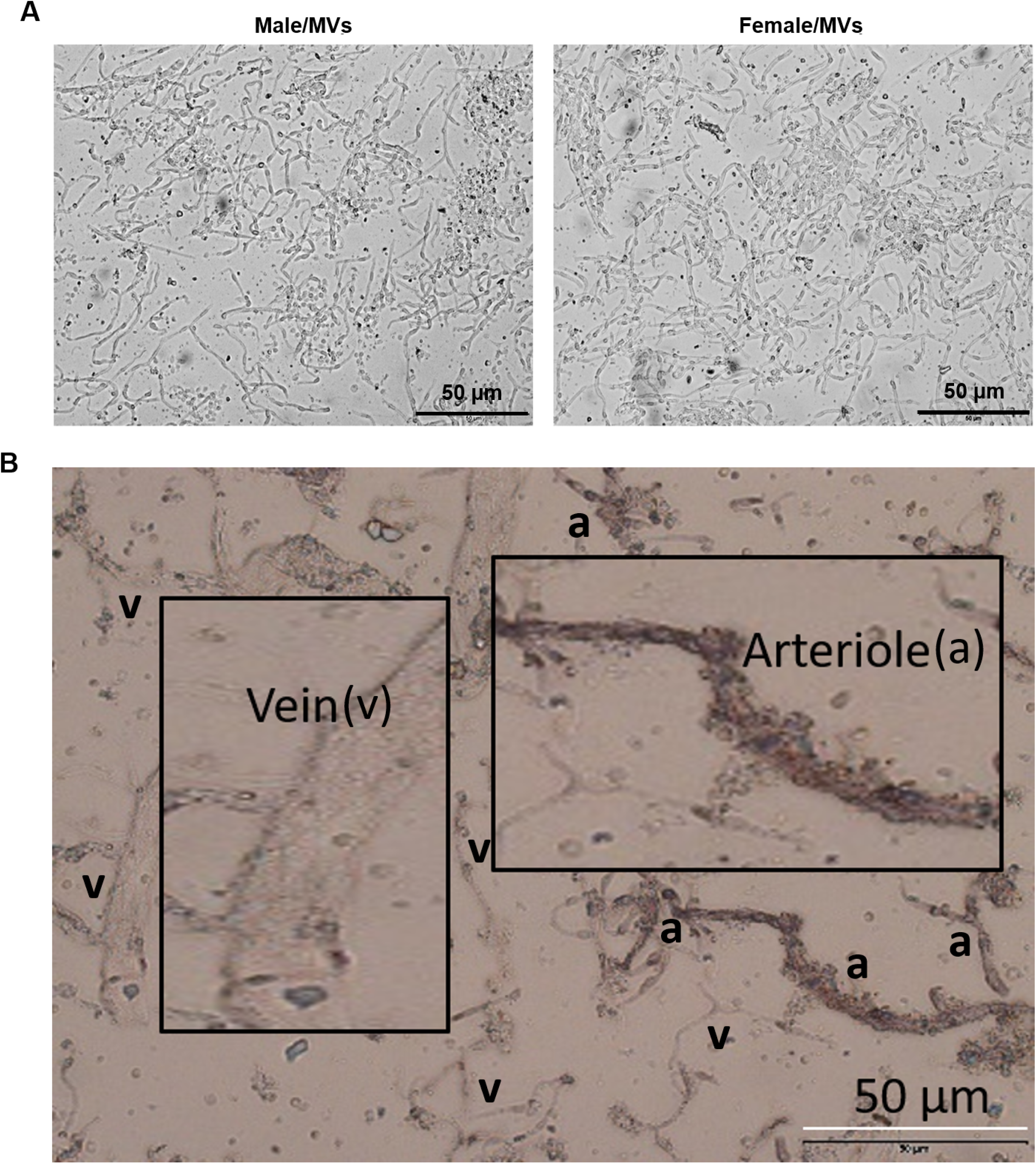
Brain MVs from male and female rats. (A) Representative images of the MVs isolated from Sprague-Dawley male (left) and female (right) rats at 20X magnification. MV preparation consisted of end arterioles, capillaries, and venules. (B) Alkaline phosphatase staining allowed discrimination between the arteriole (a) and venule (v) segments.

### MS analysis proteins in female and male rat MVs

Three technical repeats were performed for each biological sample. A discovery-based quantitative proteomics workflow using tandem mass tags (TMT) was performed. Data acquisition utilized a MS fragmentation (MS3) approach for both identification and quantitation. Proteome Discoverer 2.2 was utilized for initial data analysis. MS analysis included abundance ratios (female/male), p-values, adjusted p-values, SEQUEST-HT and PEP scores, % coverage, peptide spectral matches, and the number of peptides and unique peptides observed. Only one high scoring peptide was required for protein identification and inclusion in the results. The proteins identified in each technical repeat are provided in the supplemental data **(supplemental excel data file).** A total of 1,969 proteins were identified within this dataset, of which 1,871 were quantified. Statistical analysis was performed on these quantified proteins to determine significant differences between females and males (p < 0.05). The initial analysis identified 198 proteins in significantly (p < 0.05) different quantities. Target proteins or proteins of interests were initially identified based on fold-change [log_2_(fold change of female/male)] and - log_10_(P_adj_ value). A volcano plot was constructed (Fig. 2) to display the quantitative data **(supplemental excel data file)**. Blue stars (*) above the horizontal line represent proteins with significantly different levels (p < 0.05). Green squares (□) show protein fold changes of female/male greater than 1.5-fold. Sixty-three proteins demonstrated 1.5 or more fold change in female than male MVs. In the volcano plot, some of the most upregulated proteins in male and female MVs are circled in red and purple, respectively (Fig. 2). Notably, most of the highly upregulated proteins in both male and female MVs were related to mitochondria.

**Fig. 2:**
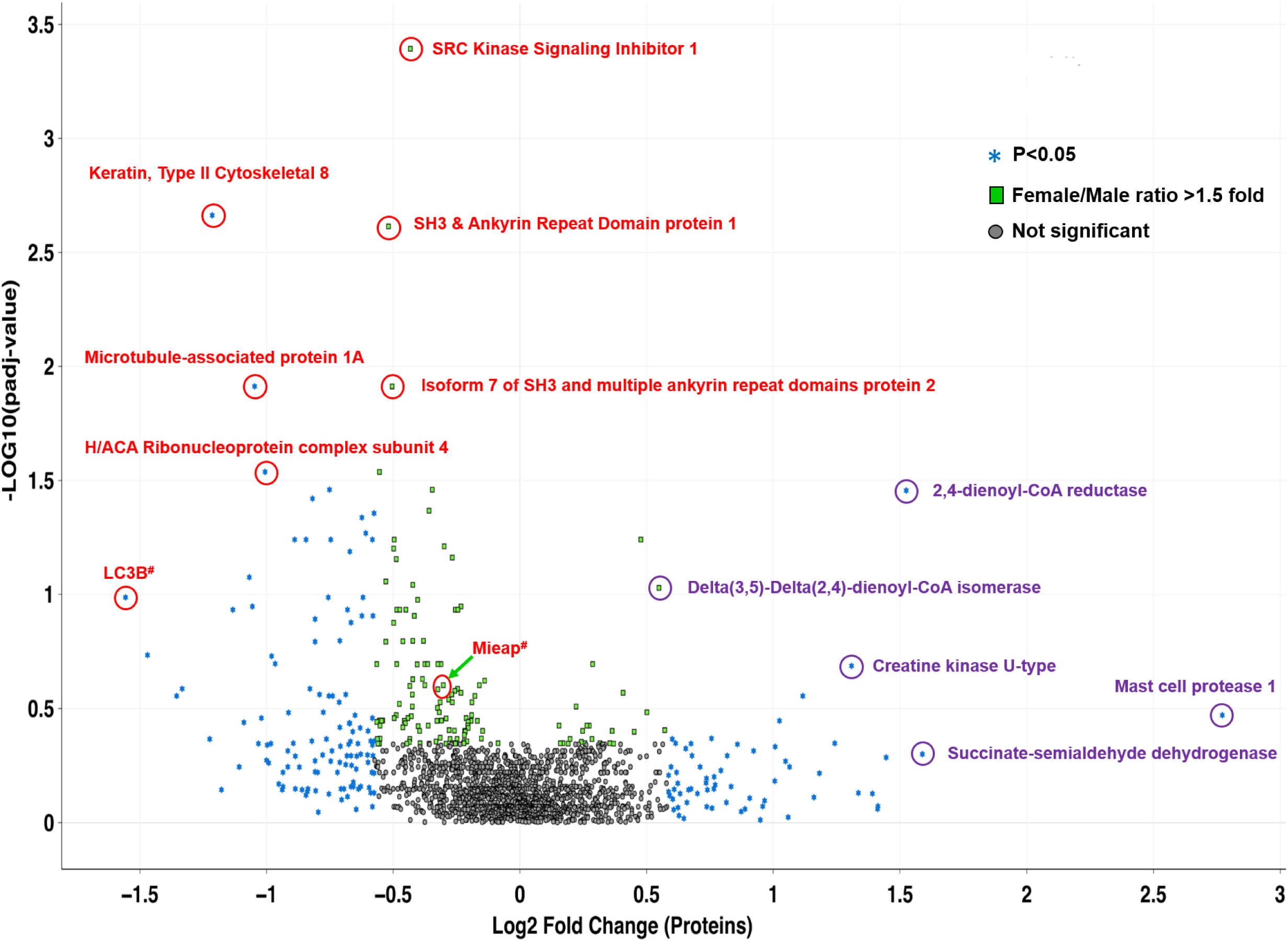
Volcano plot displaying differentially expressed proteins between female and male rat brain MVs. The vertical axis (y-axis) corresponds to the mean expression value of log10(padj-value), and the horizontal axis (x-axis) displays the log2 fold change value. Proteins of interest have been labeled and identified based on sex-based relative abundances and denoted by red and purple circles in male and female, respectively. Proteins with “green squares” have a female/male ratio of at least 1.5-fold change. Proteins identified by “blue stars” exhibited p values < 0.05. Proteins identified by the “grey dots” were quantified but did not achieve statistical significance. ^#^LC3B: Microtubule-associated proteins 1A/1B light chain 3B. ^#^MIEAP: Mitochondria eating protein.

### Classification of differentially expressed mitochondrial proteins

A total of 157 mitochondria-related proteins were quantified. There were 42 (26.7%, 42/157) significantly different (p < 0.054) mitochondria-related proteins (Fig. 3). Over half (57%; 24/42) of the mitochondrial proteins exhibited greater levels in females and nearly 43% (18/42) were more prevalent in males (Fig. 3). Similarly, 62 (39.5%, 62/157) close to significant (CS) (p ≥ 0.055 to p ≤ 0.104) mitochondria-related proteins were also analyzed (Fig. 4). Interestingly, most of the CS proteins (85.5%, 53/62) were high-abundance proteins in female MVs (Fig. 4). The mitochondrial proteins (both significant and CS) were grouped into ten categories (Fig. 5).

**Fig. 3:**
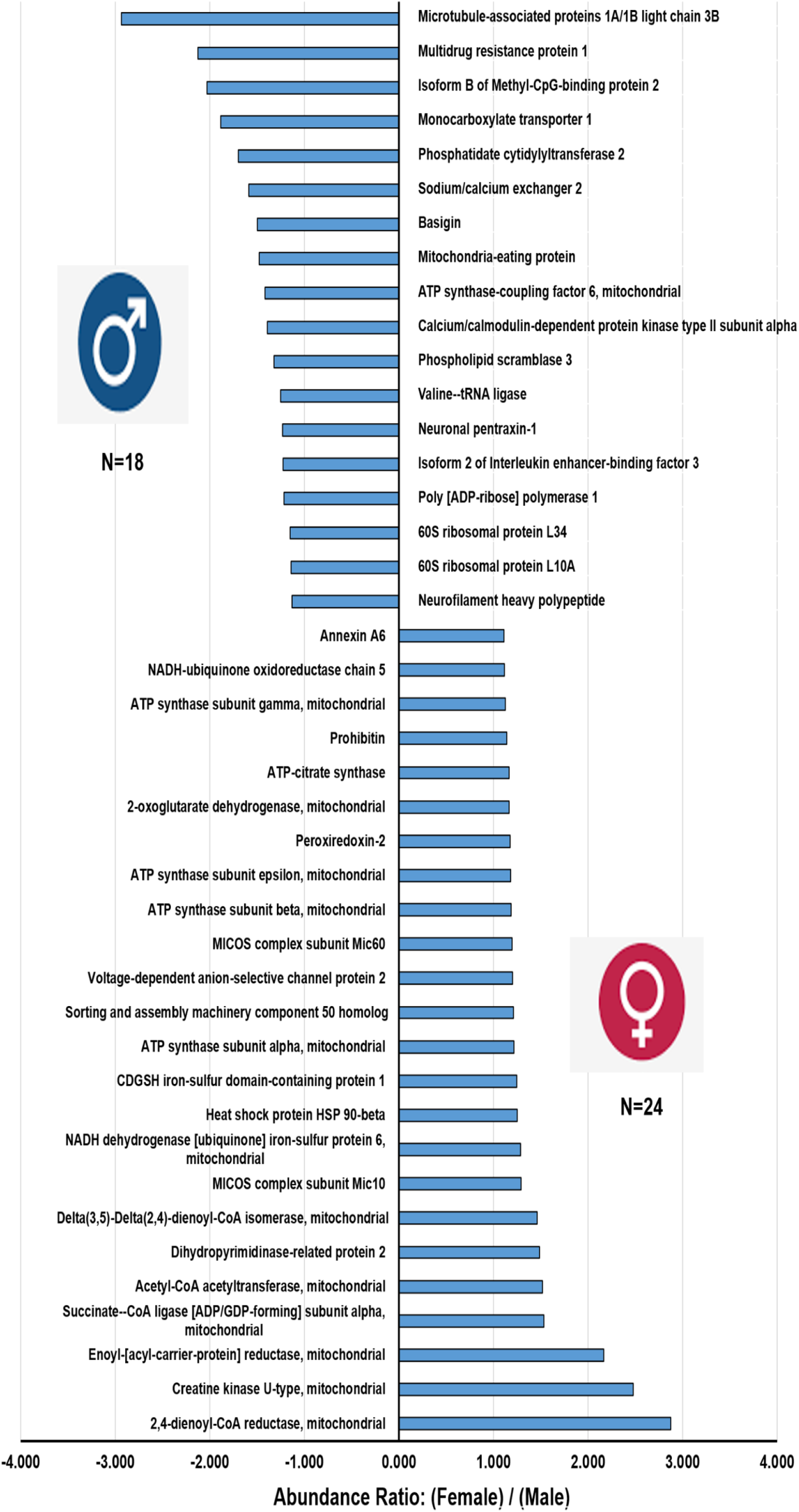
Sex dependent differences of up-regulated and down-regulated mitochondria-related proteins in brain MVs identified by TMT-based LC/MS study. The x-axis represents the abundance ratio of an average quantity of protein present in female MVs compared with male MVs (presented as Abundance ratio: Female/Male. The y-axis represents the significantly (p < 0.05) up- or down-regulated mitochondria-related proteins.

**Fig. 4:**
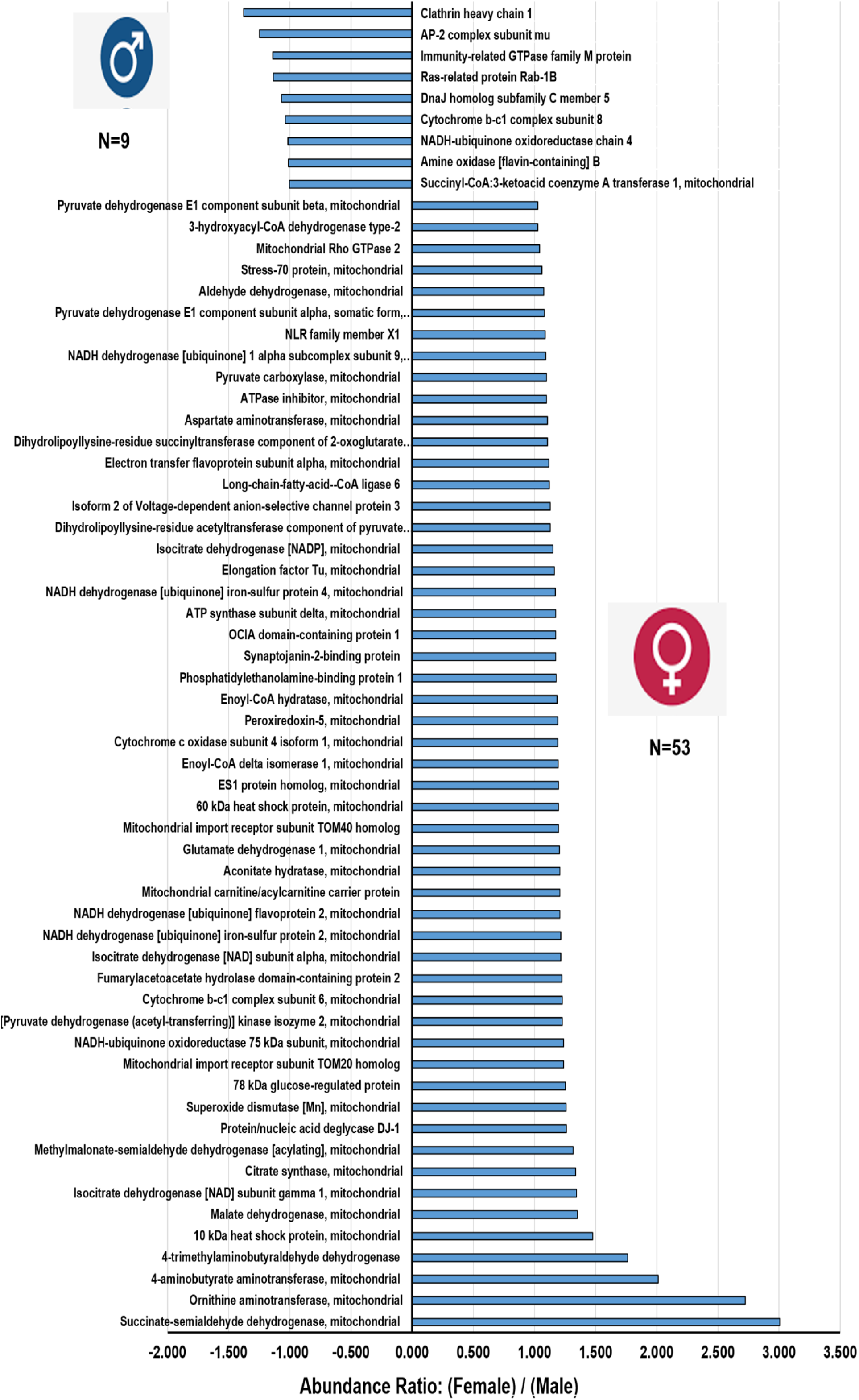
Sex-dependent differences of mitochondria-related proteins in rat microvessels approaching CS levels (p ≥ 0.055 to p ≤ 0.104). The x-axis represents the abundance ratio of an average quantity of protein present in female MVs compared with male MVs (presented as Abundance ratio: Female/Male). The y-axis represents up- or down-regulated mitochondria-related proteins.

**Fig. 5:**
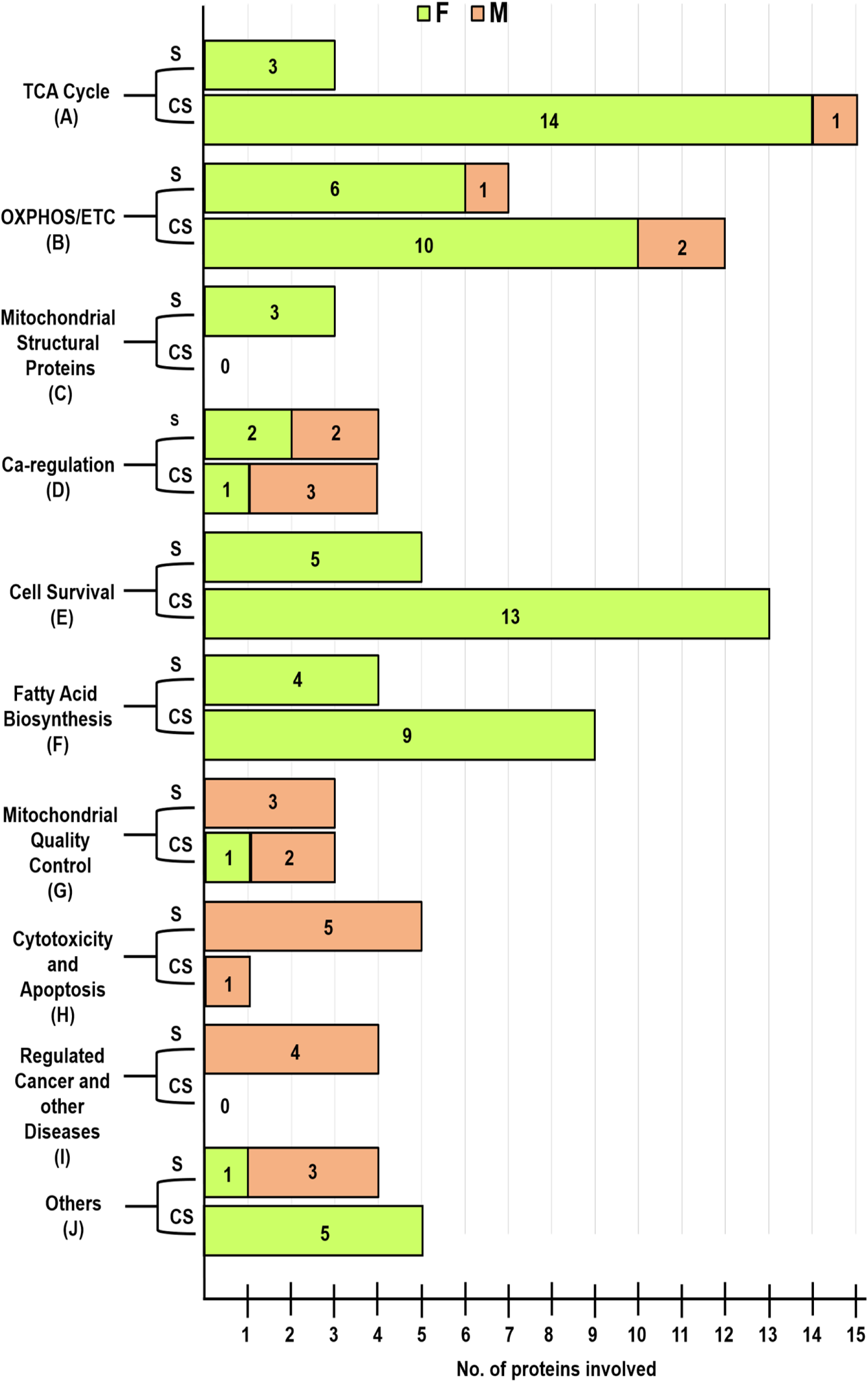
The mitochondria-related proteins were grouped into ten functional categories as shown on the y-axis. Forty-two significantly (S) different (p < 0.054) mitochondria-related proteins were clustered in different groups. Sixty-two CS (p ≥ 0.055 to p ≤ 0.104) mitochondria-related proteins were clustered in these different groups. Numbers inside the bar represent the number of proteins present in each cluster in a group.

#### Tricarboxylic acid (TCA) cycle proteins

Three high-abundance proteins were detected in female MVs: ATP-citrate synthase, 2-oxoglutamate dehydrogenase, and succinate-CoA ligase [ADP/GDP forming] subunit alpha (Fig. 3 **and** Fig. 5). Another 14 TCA cycle proteins were more prevalent in female MVs (Fig. 4 **and** Fig. 5) and those were CS (p ≥ 0.055 to p ≤ 0.104). Interestingly, no TCA cycle proteins were significantly upregulated in male MVs and only one CS protein (succinyl-CoA: 3-ketoacid coenzyme A transferase 1) was higher in male MVs (Fig. 3**‒**5).

#### OXPHOS/electron transport chain (ETC) system proteins

Seven OXPHOS/ETC system proteins were significantly (p < 0.05) upregulated and 85.7% (6/7) were observed in female MVs: ATP synthase subunit alpha, -beta, -gamma, -epsilon, NADH-ubiquinone oxidoreductase chain 5, and NADH-ubiquinone dehydrogenase iron-sulfur protein 6. The only protein that was significantly higher in male MVs was ATP synthase coupling factor 6 (Fig. 3 **and** Fig. 5). A similar trend was observed in the cluster of CS proteins. Nearly 83% (10/12) of OXPHOS/ETC system proteins were more prevalent in female than male MVs (Fig. 4 **and** Fig. 5).

#### Mitochondrial structural proteins

Three mitochondrial structural proteins were significantly higher (p < 0.05) in female vs. male MVs. MICOS complex subunit Mic10, and -Mic60 are involved in maintaining mitochondrial morphology. Sorting and assembly machinery component 50 homolog proteins participate in maintenance of mitochondrial cristae structure and assembly of the respiratory chain complex (Fig. 3 **and** Fig. 5).

#### Calcium regulation

We observed annexin A6 and voltage-dependent anion-selective channel protein 2, which were more prevalent (p < 0.05) in female MVs (Fig. 3 **and** Fig. 5). Mitochondrial Rho GTPase 2 was also elevated in female MVs (Fig. 4 **and** Fig. 5).

#### Cell survival

Eighteen cell survival related mitochondrial proteins showed greater expression in female MVs. Five of them were highly significant (p < 0.05) (Fig. 3 **and** Fig. 5) and 13 of them were CS (p ≥ 0.055 to p ≤ 0.104) (Fig. 4 **and** Fig. 5). Peroxiredoxin-2, aldehyde dehydrogenase, peroxiredoxin-5, mitochondrial import receptor subunit TOM40 homolog, superoxide dismutase (Mn), protein/nucleic deglycase DJ-1, and 4-aminobutyrate aminotransferase are all involved in protecting cells against oxidative stress and hypoxia. Heat shock proteins HSP 90-beta and 60 kDa, and 78 kDa glucose-regulated protein controlled protein folding and promoted cell survival. Prohibitin, stress-70 protein, and NRL family member X1 regulated cell proliferation and attenuated apoptosis.

#### Fatty acid biosynthesis (FAB)

Thirteen FAB-related proteins were higher in female MVs. Four were significantly elevated (p < 0.05) (Fig. 3 **and** Fig. 5): acetyl-CoA acetyltransferase, enoyl-(acyl-career-protein) reductase, 2,4-dienoyl-CoA reductase, and delta(3,5)-delta(2,4)-dienoyl-CoA isomerase. Nine were CS (Fig. 4 **and** Fig. 5): 3-hydroxyacyl-CoA dehydrogenase type 2, aspartate aminotransferase, electron transfer flavoprotein subunit alpha, long-chain-fatty-acid-CoA ligase 6, enoyl-CoA hydratase, enoyl-CoA delta isomerase 1, mitochondrial carnitine/acylcarnitine career protein, methylmalonate-semialdehyde dehydrogenase, and 4trimethylaminobutyraldehyde dehydrogenase.

#### Mitochondrial quality control

We observed six mitochondrial quality control (QC) related proteins of which five were more prevalent in male compared with female MVs. Mitochondrial eating protein (MIEAP), microtubule-associated proteins 1A/1B light chain 3B (LC3B), and monocarboxylate transporter 1 (MCT1) were significantly (p < 0.05) upregulated in male MVs (Fig. 3**-**5). Immunity-related GTPase family M protein and Ras-related protein Rab-1B were also more abundant in male MVs (Fig. 4 **and** Fig. 5).

#### Cytotoxicity and apoptosis

We observed that male MVs showed significantly higher levels of six cytotoxic and apoptosis-related proteins. Five of them (multidrug resistant protein 1, basigin, phospholipid scramblase 3, neuronal pentraxin-1, and poly[ADP-ribose] polymerase 1) were significantly (p < 0.05) greater (Fig. 3 **and** Fig. 5) and one of them (amine oxidase [flavin-containing] B) was CS (Fig. 4 **and** Fig. 5).

#### Disease related proteins

We observed four cancer or other disease-related mitochondrial proteins that showed high expression (p < 0.05) in male MVs (Fig. 3 **and** Fig. 5). Neurofilament heavy peptide, 60S ribosomal protein L34, Isoform B of methyl-CpG-binding protein 2, and 60S ribosomal protein L10A were all abundant in male MVs.

#### Others

A total of nine proteins (four significant, and five CS), which were not discussed in the previously mentioned categories, showed sex differences. Phosphatidate cytidylyltransferase 2 (involved in glycerophospholipid metabolism), valine-tRNA ligase (involved in tRNA biosynthesis), and isoform 2 of interleukin enhancer-binding factor 3 (participating in transcription/post-transcription) were more abundant (p < 0.05) in male MVs. CDGSH iron-sulfur domain-containing protein 1 (involved in adipogenesis) was more highly expressed (p < 0 .05) in female MVs (Fig. 3 **and** Fig. 5). Conversely, OCIA domain-containing protein 1 (participating in stem cell differentiation), synaptojanin-2-binding protein (regulating endocytosis), phosphatidylethanolamine-binding protein 1 (modulating signalling pathways), isoform 2 of voltage-dependent anion-selective channel protein 3 (participating in energy metabolism), and ornithine aminotransferase (involved in the formation of proline) were highly abundant (CS) in female MVs (Fig. 4 **and** Fig. 5).

### Western blots of key proteins

Three proteins showing prominent male/female differences and relevant to our conclusions concerning mitochondrial degradation or FFA metabolism were selected for further validation. Similar to the Proteomics results, we found with western blotting that levels of LC3B and MCT1 were higher in male than female MVs (p<0.05) and that DECR1 was more abundant in female than male MVs (p<0.05) (Fig. 6).

**Fig. 6:**
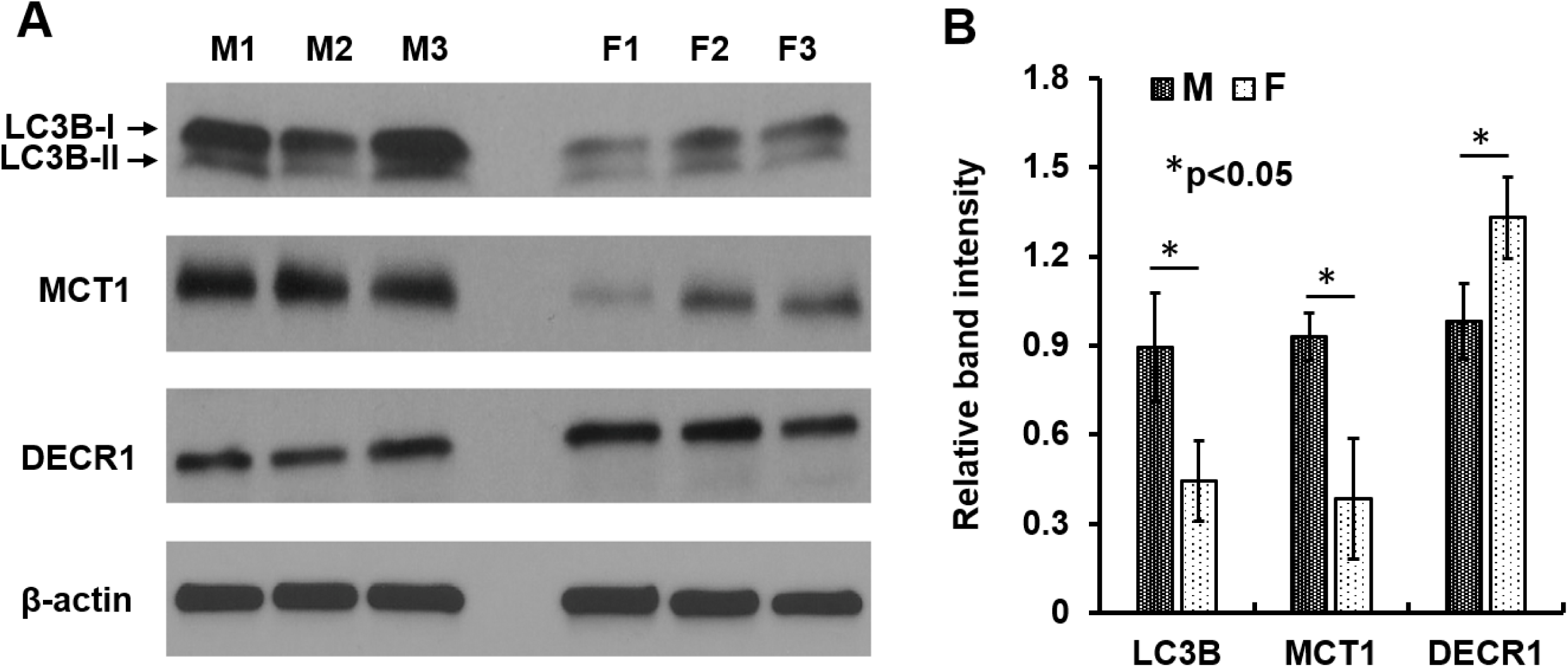
Western blots were performed to validate the LC-MS/MS-based proteomics data. (A) MVs form age-matched three males (M1, M2, and M3) and three females (F1, F2, and F3) were isolated at the same time, lysed the MVs, and equal amount (20 µg) of proteins were loaded in the gels and the said proteins were detected by immunoblots. (B) The relative band intensity was compared by the Image-J software (version 1.50). Values are mean±SEM and significant changes are presented as *p<0.05. LC3B: Microtubule-associated proteins 1A/1B light chain 3B; MCT1: Monocarboxylate transporter 1; DECR1: 2,4-dienoyl-CoA reductase.

### Top canonical pathways are linked with mitochondria

Included in **Table 1** are the top five canonical pathways with coverage and confidence scores, as well as the top five associated network functions. The top five canonical pathways that exhibit sex-based differences include mitochondrial dysfunction, oxidative phosphorylation, sirtuin signaling, TCA cycle, and eIF2 signaling. Figure 7 illustrates the top overlapping canonical pathways in rat brain MVs. These were identified based on the coverage and overlap of the number of proteins known to be associated with the described pathways, as well as the confidence of the peptide false discovery rate (FDR) (< 1%). Each signaling pathway within this figure shares a minimum of 10 “over-lapping” proteins with one or more neighboring nodes, representing respective signaling pathways. The subsequent p-values of overlap are also displayed within this figure. In Figure 7, we present the top 31 canonical pathways exhibiting sexual disparity in female and male MVs. Most of the top overlapping canonical pathways were either directly or indirectly linked with mitochondria. Oxidative phosphorylation, TCA cycle, fatty acid β-oxidation (FAO), and calcium signaling pathways are exclusively related to mitochondria.

**Fig. 7:**
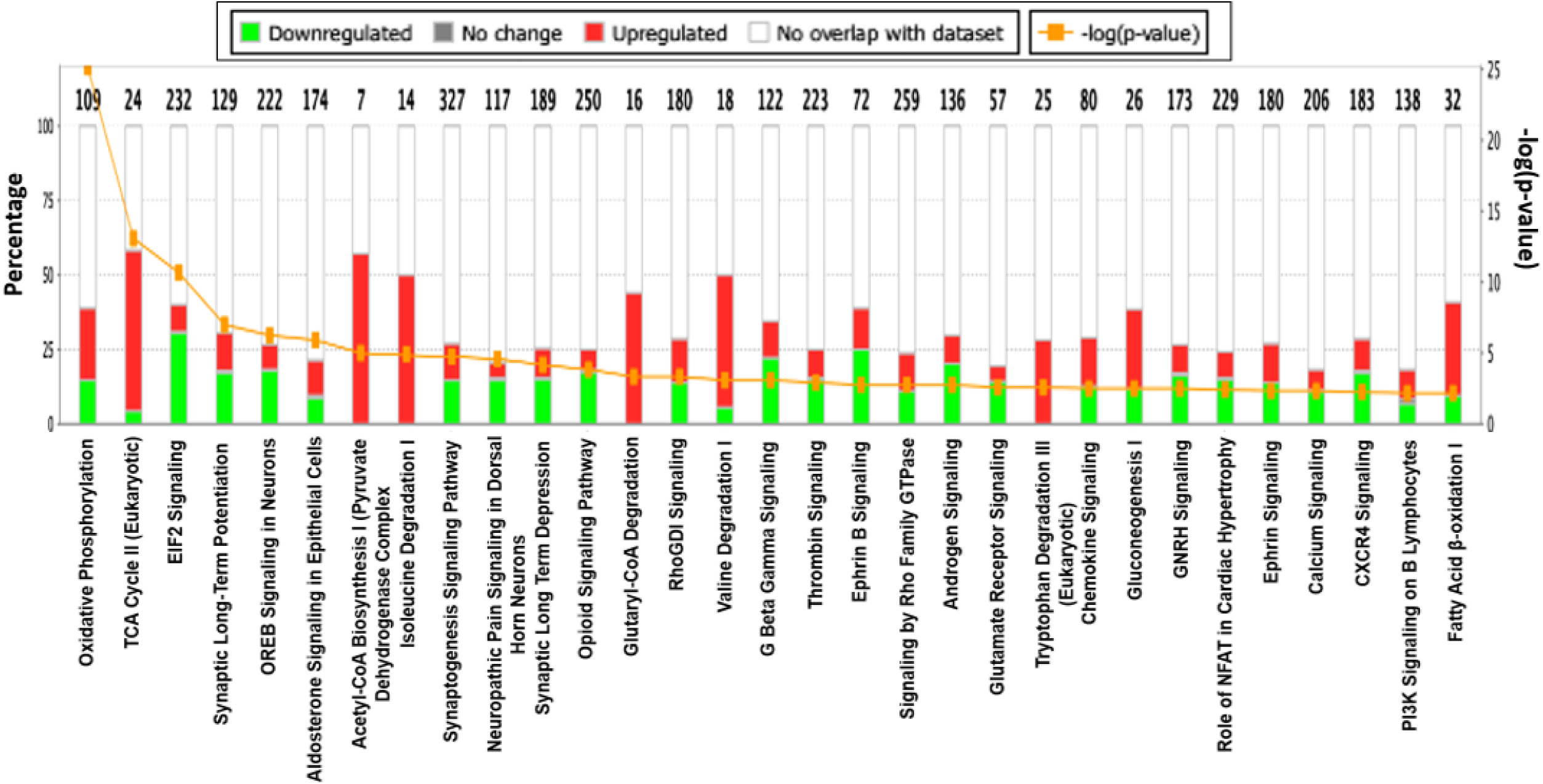
Top canonical pathways exhibiting sexual disparity in MVs between male and female rats. This figure is based on proteins identified in the dataset and their known (published) regulatory associations within the listed pathways (x-axis). Bar graphs illustrate the total number of proteins in each pathway, and the number of genes differentially expressed in each pathway, indicated on the top of each bar (y-axis). The MS comparison was defined to describe female/male ratios. Green color indicates proteins more abundant in male rat microvessels whereas red color indicates proteins more prevalent in females. The orange [-log(p-value)] line illustrates the “p-value of overlap” of the proteins in our dataset relative to IPA’s predefined canonicals. Each histogram bar can be expanded into larger canonical signaling networks within the software for further details, including protein-protein interactions (both upstream and downstream from a selected node).

**Table 1:** Summary report generated using Ingenuity Pathway Analysis depicting the top 5 canonical pathways enriched in the data set with confidence (p-value of overlap) and coverage (overlap of proteins) of total pathway. Additionally, the top 5 toxicology lists for this dataset are listed below the table.

|  |  |  |
| --- | --- | --- |
| Mitochondrial Dysfunction | 1.32E-33 | 26.3% (45/171) |
| Oxidative Phosphorylation | 7.37E-26 | 29.4% (32/109) |
| Sirtuin Signaling Pathway | 6.06E-18 | 13.4% (39/292) |
| TCA Cycle II (Eukaryotic) | 8.51E-14 | 50.0% (12/24) |
| EIF2 Signaling | 2.20E-11 | 11.6% (27/232) |
| **<u>Top 5 Toxicology List</u>** |  |  |
| 1. Mitochondrial Dysfunction |  |  |
| 2. Fatty Acid Metabolism |  |  |
| 3. Cardiac Necrosis/Cell Death |  |  |
| 4. Decreased Permeability of Mitochondria and Mitochondrial Membrane |  |  |
| 5. Renal Necrosis/Cell Death |  |  |

**Table 1:** Summary report generated using IPA. Table depicts the top 5 canonical pathways enriched in the data set with confidence (p-value of overlap) and coverage (overlap of proteins) of total pathway. The top 5 toxicology lists for this dataset are listed additionally below the table.
| **<u>Name</u>** | **<u>p-value</u>** | **<u>Overlap</u>** |
| --- | --- | --- |
| Mitochondrial Dysfunction | 1.32E-33 | 26.3% (45/171) |
| Oxidative Phosphorylation | 7.37E-26 | 29.4% (32/109) |
| Sirtuin Signaling Pathway | 6.06E-18 | 13.4% (39/292) |
| TCA Cycle II (Eukaryotic) | 8.51E-14 | 50.0% (12/24) |
| EIF2 Signaling | 2.20E-11 | 11.6% (27/232) |
| **<u>Top 5 Toxicology List</u>** |  |  |
| 1. Mitochondrial Dysfunction |  |  |
| 2. Fatty Acid Metabolism |  |  |
| 3. Cardiac Necrosis/Cell Death |  |  |
| 4. Decreased Permeability of Mitochondria and Mitochondrial Membrane |  |  |
| 5. Renal Necrosis/Cell Death |  |  |

Acetyl-CoA biosynthesis and isoleucine/valine degradation pathways are associated with the TCA cycle, glucogenesis, tryptophan degradation, glutamate receptor signaling, G beta gamma signaling, and Rho family GTPase pathways all spanned both cytoplasmic and mitochondrial compartments.

## DISCUSSION

### Overview

The major finding of this study is that substantial sex-dependent changes in energy producing proteins involving the TCA cycle, OXPHOS, and FAO are present in cerebral MVs of young, adult rats. These results confirm and extend previous studies performed on large cerebral arteries showing that mitochondrial proteins (proteins in complexes I-V, voltage-dependent anion channel proteins, etc.), as well as mitochondrial DNA, are more highly expressed in female than male rats (Rutkai et al, 2015; Lee et al, 2019b). Elevated levels of mitochondrial proteins in large, female arteries was associated with greater mitochondrial respiration and enhanced mitochondrial-mediated dilation of arteries compared with males (Rutkai et al, 2015; Demarest et al, 2016). We have extended these earlier findings with a more comprehensive Proteomics examination of proteins in cerebral MVs, with an emphasis not only on individual mitochondrial structural proteins and energy producing proteins but also on proteins known to modulate mitochondrial numbers and function. We studied cerebral MVs because of recent recognition that this vascular segment is more susceptible to dysfunction during aging and disease than large arteries and that MVs are a primary initiation site for development of severe neurological diseases such cognitive impairment, vascular dementia, and AD (Liu et al, 2018; Bourassa et al, 2019; Erdő & Krajcsi 2019). Although several current findings confirmed results of our earlier studies, the expansive and thorough nature of the present study has yielded novel and surprising findings. These findings include extensive information regarding the numerous proteins involved in the TCA cycle and OXPHOS, the balance between mitochondrial protective and destructive proteins, and information about proteins that facilitate the production of ATP via the FAO pathway. Western blotting of proteins involved in mitochondrial degradation or the FAO pathway validated our key Proteomics results. Overall, the results promote the view that female cerebral MVs have greater energy generating capacity via mitochondrial dependent pathways, are more resistant to mitochondrial stress and destruction, and have greater ability to use fuels alternative to glucose such as fatty acids. Furthermore, major sex differences extend to proteins not directly involved in energy production or modulation of mitochondrial function, including calcium regulation and cellular signaling.

### Energy producing proteins

Three main enzymatic pathways (TCA cycle, OXPHOS, and FAO) leading to ATP production are more prominent in female than male cerebral MVs. However, we could not detect a consistent difference between male and female cerebral MVs in proteins involved in glycolysis (data not shown). Previous studies using more traditional approaches on large cerebral arteries have shown that mitochondrial proteins and mitochondrial-related mechanisms under baseline conditions are more highly expressed in female than male rats. Thus, using Western blotting, proteins located in the inner and outer membranes are present in high concentrations in female compared with male arteries (Rutkai et al, 2015). These proteins were encoded by both nuclear and mitochondrial DNA, indicating a coordinated synthesis and assembly system (Burstein et al, 2018). Our previous findings that mitochondrial respiration and mitochondrial-induced signaling are enhanced in female arteries are consistent with these results (Rutkai et al, 2015). As expected, proteins such as MICOS complex subunit 60, which plays a crucial role in maintenance of crista morphology, is also expressed more in female cerebral MVs. We took a “robust” approach in this survey of MV proteins by considering, in some analyses, sex differences falling within the CS range (p > 0.05 to p < 0.10) as well as the traditional p < 0.05 designation. We did this to allow a broad picture of the information generated by the proteomics approach and to guide future studies. Extrapolating from our results on large arteries, we expected that mitochondrial respiration and the production of biologically important mitopeptides to be enhanced in female compared with male cerebral MVs.

Another novel finding of this study is that FAO proteins are enriched in female cerebral MVs. 2,4-dienoyl-CoA reductase and Delta(3,5)-Delta(2,4)-dienoyl-CoA isomerase both were highly abundant in female MVs and were involved in fatty acid metabolism. While the traditional view posits that the brain, encompassing neurons, glia cells, and blood vessels utilizes glucose exclusively as an energy source, findings from astroglia challenge this view. Recent research shows that up to 20% of the energy production in the brain arises from fatty acids (Romano et al, 2017; Snowden et al, 2017). Thus, the brain uses an energy-efficient process where the complete oxidation of a 16-carbon palmitic acid yields 112 molecules of ATP. Increased levels of FAO are crucial for homoeostatic regulation, specifically during fasting or endurance exercise that requires high-energy production.

### Mitochondrial quality control (QC)

One of the most striking findings of this study was the greater abundance in male cerebral MVs of proteins involved in the degradation of mitochondria. These included microtubule-associated 1A/1B light chain 3B (LC3B) (Fig. 3), which contributes to mitochondrial quality and quantity by eliminating damaged or dysfunctional mitochondria. Mitochondrial eating protein (MIEAP) (Fig. 3), a key p53-inducible protein, which mediates the repair or degradation of unhealthy mitochondria, and phospholipid scramblase 3, which promotes translocation of cardiolipin from the inner to outer mitochondrial membrane and leads to mitophagy are other examples of mitochondrial QC. MIEAP also plays a pivotal role in mitochondrial QC by repairing or eliminating unhealthy mitochondria via MALM (<u>M</u>IEAP-induced <u>a</u>ccumulation of lysosome-like organelles within <u>m</u>itochondria) or MIV (<u>M</u>IEAP-induced <u>v</u>acuole) generation, respectively (Kitamura et al, 2011; Nakamura & Arakawa 2017). Damaged mitochondria can be discarded by an organelle specific form of autophagy known as mitophagy or mitochondrial autophagy. A critical protein, LC3B, localizes to the autophagosome membrane and thus is the key regulator involved in autophagosome formation and is widely used as a marker of autophagic activity (Kabeya et al, 2000; Mizushima et al, 1998; Mizushima et al, 2001). Aberrant expression of LC3B has been reported in several solid tumors (Giatromanolaki et al, 2014; Huang et al, 2010; Ko et al, 2013; Lazova et al, 2012) and promotes translocation of cardiolipin from the inner to the outer mitochondrial membrane, which promotes BID recruitment and enhances tBid-induced mitochondrial damage. PLS3(F258V)-overexpressing cells have decreased mitochondrial mass shown by lower cytochrome c and cardiolipin content, poor mitochondrial respiration, and reduced oxygen consumption and intracellular ATP; whereas, wild-type PLS3-transfected cells exhibit increased mitochondrial mass and enhanced respiration (Liu et al, 2003; Kowalczyk et al, 2009). Although an optimal balance between mitogenesis and mitophagy is critical to the health of the vascular cells according to physiological status, it appears that mechanisms achieving this balance are different in males and females.

### Canonical pathways

The most prominent canonical pathways revealed by proteomic analysis were either directly or indirectly linked with mitochondria. Oxidative phosphorylation, the TCA cycle, FAO, are exclusively dependent on mitochondria, and calcium signaling pathways in mitochondria are important to maintain cell homeostasis. Acetyl-CoA biosynthesis and isoleucine/valine degradation pathways were associated with the TCA cycle. Glucogenesis, tryptophan degradation, glutamate receptor signaling, G beta gamma signaling, and Rho family GTPase pathways all span both cytoplasmic and mitochondrial compartments. Annexin A6 regulate mitochondrial morphogenesis and mitochondria lacking annexin A6 are fragmented and exhibit impaired respiration and reduced mitochondrial membrane potential (Chlystun et al, 2013; Banerjee et al, 2015). Mitochondrial Rho GTPase (MIRO) is a key regulator of mitochondrial transport and dynamics and is involved in diseases associated with defects in mitochondrial movement and function, especially neurodevelopmental and neurodegenerative disorders (Tang 2015). Overexpression of MIRO with other interactors prevents amyloid β (Aβ)-induced mitochondrial fragmentation (Werle et al, 2014). MIRO1 is significantly reduced in the spinal cord tissue of amyotrophic lateral sclerosis (ALS) patients (Zhang et al, 2015). On the other hand, sodium/calcium exchanger 2 and calcium/calmodulin-dependent protein kinase type II subunit alpha proteins are abundantly higher (p < 0.05) in our male MVs. Cytosolic and mitochondrial Ca^2+^ overload are principal pathophysiological events during brain ischemia. Most studies suggest a protective effect on cells of blocking the mitochondrial exchanger activity (Castaldo et al, 2009). Clathrin heavy chain 1, AP-2 complex subunit mu, and DnaJ homolog subfamily C member 5 proteins were also highly abundant (CS) in male MVs. Thus, mitochondrial-related associations account for the majority of sex-related differences in proteins in rat MVs.

### Physiological significance of sex differences

The underlying basis for sex differences in the cerebral circulation of rats appears to be estrogen. Several studies have shown that estrogen increases mitochondrial numbers and efficiency in cerebral endothelium and increases protection against ROS via regulation of peroxisome proliferator-activated receptor-γ coactivator-1α (Kemper et al. 2013, 2014). Although female cerebral MVs have greater energy producing capability via the TCA and OXPHOS pathways and enhanced ability to use fatty acids via FAO, the physiological importance of this finding is unclear. One possible explanation is that enhanced energy production capacity and flexibility of fuel source represents a generalized phenotype occurring in all regional circulations and is preparatory to the morphological and physiological changes to the microvasculature that occur during menstruation, pregnancy, and lactation. Obvious and widespread vascular and metabolic changes occur during pregnancy, post-partum, and during lactation. For example, a switch from anabolic to catabolic metabolism occurs throughout pregnancy and thus the ability of the cells, including the vasculature, to use free fatty acids as well as glucose becomes important. Major changes in metabolism also occur during lactation. There are major changes in the cardiovascular system associated with menstruation, pregnancy, and lactation such as substantial changes in plasma volume and circulating factors. With respect to the cerebral circulation, changes in neurovascular coupling occur during the estrus cycle (Capone et al, 2009) and changes in parenchymal blood vessels and blood-brain barrier function occur during pregnancy (Johnson & Cipolla 2015). For example, Cipolla and colleagues have reported that brain MVs undergo outward hypotrophic remodeling and capillary density in the posterior cerebral cortex increases during pregnancy (Johnson & Cipolla 2015). Greater mitochondrial numbers in females may also lead to enhanced secretion of beneficial mitopeptides which protect cells against stress. Mitopeptides can act within a cell or be excreted into the extracellular fluid to act on adjacent or distant cells (Bachar et al, 2010;Yen et al, 2013).

### Limitations of the study

There are several limitations of our study. First, we took a “robust” approach and considered proteins that reached the p < 0.05 level as well as those that were close to significant (CS): p > 0.05 to p < 0.10. We did so to fully maximize the usefulness of the data generated by a proteomics approach and to plan future studies. Second, our MV preparation contained different segments of the microvasculature and did not represent solely arterioles, capillaries, or venules. Thus, we cannot attribute our results to any one segment or cell type of the microvasculature. However, we are unaware of procedures that would allow us to separate vascular segments with any degree of confidence and still yield enough protein for our analyses. Third, we did not evaluate the activity of the proteins of interest. However, our prior studies have shown that mitochondrial numbers, mitochondrial mediated signaling, and mitochondrial respiration are greater in female than male large cerebral arteries in rats. And fourth, a substantial amount of information provided by our proteomics analysis was excluded from the discussion because of lack of perceived relevance to the current focus on mitochondria. However, future studies might establish important but previously unrecognized links to mitochondria that further inform MV sexual dimorphism and its clinical implications. In conclusion, we provide the first comprehensive examination of mitochondria-related proteins in male and female MVs of rats and show that female mitochondria are more numerous, stable, and versatile than male mitochondria.

## MATERIALS AND METHODS

### Animals

Age-matched, male and female, Sprague-Dawley rats (10–12 week old, n = 6) were obtained from Charles River Laboratories (Wilmington, MA, USA) and housed according to the Institutional Animal Care and Use Committee of Tulane University guidelines, and were also in compliance with National Institutes of Health Office of Laboratory Animal Welfare guidelines. The Institutional Animal Care and Use Committee of Tulane University approved the protocols, which were also in compliance with ARRIVE (Animal Research: Reporting in Vivo Experiments) guidelines. Rats had free access to food and water, ad libitum, with a standard light/dark cycle.

### Microvessels isolation

The MV isolation procedure from rat brains was modified from previously published protocols (Silbergeld & Ali-Osman 1991; Yousif et al, 2007; Merdzo et al. 2016, 2017; Sure et al, 2018). In brief, the rat brain was removed and placed on filter paper and the hindbrain, olfactory lobes, white matter, and large surface blood vessels were removed and discarded. The remaining cortical tissue was homogenized with 5 mL ice cold Dulbecco’s phosphate buffered saline (DPBS) (Life Technologies Corporation, NY, USA) on ice. The homogenate was placed into a new 50 mL conical tube, centrifuged at 3300 × g for 15 min, the supernatant was carefully removed, and the pellet was resuspended in 10 mL of 17.5% dextran. The suspension was passed through a 300 μm filter (pluriSelect Life Science, CA, US) using a 5 mL pipette to remove unhomogenized tissue pieces. The filtrate was centrifuged at 7900 × g for 15 min and contaminated myelin was removed. The MV pellet was gently resuspended in 2% BSA and passed through a 70 μm filter (Falcon, Corning Incorporated, NY, USA). The MVs were placed into a 1.5 mL tube, centrifuged at 13.2 × g for 15 min, and a final clean-up of any remaining myelin was done using 17.5% dextran followed by 2% BSA. The MVs were resuspended in 100 µL of DPBS and stored at −80°C until further use.

### Alkaline Phosphatase (AP) staining

An AP staining protocol was used to differentiate between arterioles, capillaries, and venules and was adopted from a published protocol (Eliceiri et al, 2011) with some modification. An aliquot of 10‒15 µL from 100 µL of MVs was placed into a 1.5 mL tube and washed in developing buffer (0.1M Tris-HCL, pH 9.5 + 0.1M Nacl + 0.05M Mgcl_2_) for 10 min. The preparation was centrifuged at 7900 × g for 15 min and incubated with nitro blue tetrazolium (NBT)/5-bromo-4-chlori-3-indoyi-phosphate (BCIP) substrate solution. An NBT/BCIP stock solution was prepared by adding 18.75 mg/mL of NBT (Boehringer Mannheim GmbH, Germany) + 9.4 mg/mL of BCIP (Boehringer Mannheim GmbH, Germany) in 70% dimethyl sulfoxide (Sigma-Aldrich Co., St Louis, MO). To prepare the NBT/BCIP substrate solution, 200 µL of NBT/BCIP stock solution was added to 9.8 mL developing buffer. The color development was monitored under light microscopy and representative pictures were taken with 20X magnification.

### Western blot analysis

Protein lysates from MVs were prepared as described previously (Merdzo et al, 2017; Rutkai et al, 2015). Equal amount (20 µg) of samples were separated by gel electrophoresis on a 4-20% SDS-PAGE gradient gel, and then transferred onto a PVDF membrane, blocked with casein blocking buffer (Li-cor, Lincoln, NE), and incubated overnight with primary antibodies. The following primary antibodies were used: anti-Microtubule-associated proteins 1A/1B light chain 3B (LC3B-I/-II) at 1:1000 dilution (14, 16 kDa, Cell Signaling Technology, Danvers, MA); anti-monocarboxylate transporter 1 (MCT1) at 1:1000 dilution (54 kDa, EMD Millipore Corporation, Temecula, CA), anti-2,4-dienoyl CoA reductase (DCAR1) at 1:1000 dilution (36 kDa, Novus Biologicals, Centennial, CO). The internal control was β-actin at 1:5,000 dilution (42 kDa; No. A5441, Sigma-Aldrich, St. Louis, MO). After incubation with the primary antibody, the membranes were washed and incubated with either secondary goat anti-rabbit IgG or goat anti-mouse IgG at 1:5,000 dilution (Cell Signaling Technology, Danvers, MA) at room temperature for 90 min. Immunoblots were visualized using chemiluminescence (LumiGLO, Gaithersburg, MD) and autoradiography. The optical density of the specific bands was quantified, normalized to the intensity of the corresponding β-actin band using ImageJ software (version 1.50) and compared.

### Proteomics Sample Preparation

The samples were prepared with a lysis buffer containing 1% SDS with intermediate vortexing and sonication. The lysate was stored at −80°C until further use. Protein quantitation was performed using bicinchoninic acid (BCA) Protein Assay Kit (Thermo Scientific, Rockford, IL); 100 µg of each sample was used for TMT labeling.

### Discovery-based quantitative proteomics

Each sample was prepared for trypsin digestion by reducing the cysteines with tris(2-carboxyethyl) phosphine (TCEP) followed by alkylation with iodoacetamide (IAA). After chloroform-methanol precipitation, each protein pellet was digested with 2 µg sequencing grade trypsin (Thermo Scientific, Rockford, IL, USA) overnight at 37°C. Proteolytic peptides were labeled using a TMT 6-plex reagent kit (Thermo Scientific) according to the manufacturer’s protocol. An equal amount of each TMT-labelled sample was mixed in a single tube with Sep-Pak purified (Waters, Ireland) using acidic reverse phase conditions. After drying to completion, off-line fractionation was employed to reduce the sample complexity. The pH of the samples was adjusted with 20 mM ammonium hydroxide (AH). This mixture was subjected to basic pH reverse phase chromatography (Dionex U3000, ThermoFisher Scientific) on an ACUITY UPLC peptide BEH C18 column, 300A, 1.7 μm, 2.1 x 50 mm (Waters, Ireland). Eluate was UV monitored at 215 nm for an injection of 100 µL at 0.1 mL/min with a gradient developed from 10 mM AH, pH 10 to 100% ACN in AH pH 10 over 90 min.

A total of 48 fractions (200 µL each) were collected in a 96-well microplate and recombined in a checkerboard fashion to create 12 “super fractions” (original fractions 1, 13, 25, and 37 became new super fraction #1, original fractions 2, 14, 26, and 38 became new super fraction #2, etc.). Each of the 12 “super fractions” were then run on a Dionex U3000 nano flow system coupled to an Orbitrap Fusion Tribrid mass spectrometer (ThermoFisher Scientific). Each fraction was subjected to on-line 90-min chromatography using a gradient from 2‒25% acetonitrile in 0.1% formic acid (ACN/FA) over the course of 65 min, a gradient to 50% ACN/FA for an additional 10-min, a step to 90% ACN/FA for 5 min and a 10 min re-equilibration to 2% ACN/FA. Chromatography was carried out in a “trap-and-load” format using a PicoChip source (New Objective, Woburn, MA); trap column C18 PepMap 100, 5 µm, 100A and the separation column was PicoChip REPROSIL-Pur C18-AQ, 3 µm, 120A, 105 mm. The entire run had a 0.3 µL/min flow rate. Electrospray was achieved at 2.6 kV.

TMT data acquisition utilized an MS3 approach for data collection. Survey scans (MS1) were performed in the Orbitrap utilizing a resolution of 120,000. Data dependent MS2 scans were performed in the linear ion trap using a collision induced dissociation of 25%. Ions were fragmented using high energy collision dissociation of 65% and detected in the orbitrap at a resolution of 30,000. This was repeated for a total of three technical replicates. TMT data analysis was performed using Proteome Discoverer 2.2. The three runs of 12 “super fractions” were merged and searched using SEQUEST. The protein FASTA database was Rattus norvegicus FASTA ID= SwissProt tax ID=10116, version 2017-10-25. Static modifications included TMT reagents on lysine and N-terminus (+229.163), carbamidomethyl on cysteines (+57.021), dynamic modifications included phosphorylation of serine, threonine, and tyrosine (+79.966 Da), and dynamic modification of oxidation of methionine (+15.9949). Parent ion tolerance was 10 ppm, fragment mass tolerance was 0.6 Da, and the maximum number of missed cleavages was set to 2. Only high scoring peptides were considered utilizing an FDR of 1%.

### Data Analyses and Bioinformatics

Initial data analysis was performed with Proteome Discoverer 2.2. Factors included in MS analysis for proteins were abundance ratios (female/male), p-values, adjusted p-values, SEQUEST-HT and PEP scores, % coverage, peptide spectral matches, the number of peptides, and unique peptides observed. Only one unique high scoring peptide was required for inclusion of protein identification in the results. Subsequent bioinformatic analyses were performed using GeneGo (PANTHER) and STRING prior to loading into Qiagen’s IPA. Following completion of data acquisition and analysis in Thermo and GeneGo, our data set was uploaded for bioinformatics using IPA. Multiple bioinformatic platforms were used to cross-reference proteins and resolve inconsistencies that may arise due to variable protein identifications across different libraries, incorrect classifications, or scheduled update differences between software packages. IPA content information: Report Date: 2019-05-31, Report Version: 485480, Content Version: 47547484 (Release Date: 2019-02-08). All confidently identified peptides, including proteins identified by one unique peptide, were exported to Microsoft Excel. When annotating and applying filters for the data set uploaded to IPA, the following criteria were applied: Only < 1% FDR SEQUEST-HT scoring peptides were analyzed: only high experimentally observed molecules were factored; those with a minimum abundance ratio ≥ 1.5 and a statistically significant p-value of ≤ 0.05 were considered in the high confidence analysis. IPA organism settings included human, mouse, and rat. These filters were applied in a “core-analysis” of the entire dataset. The initial dataset of 1,969 proteins was uploaded, filtered down to the top 20% of statistical confidence, and all proteins were differentially expressed by at least 1.5-fold change (FC), which yielded 554 “analysis ready” molecules that were used for protein enrichment analysis of the dataset. The bioinformatics reported here are descriptive of the entire 3 vs. 3 multiplexed dataset. The core analysis summary report consisted of the top canonical pathways, upstream regulator, signaling networks, and the disease, molecular, and biological functions of the MS-derived cerebral microvessel proteome of the Sprague-Dawley rat.

### Statistical analysis

Results were expressed as mean ± SEM and the number of independent measurements is indicated by ‘‘n’’. Data comparison was performed using unpaired Student’s t-test and one-way ANOVA with Tukey’s post hoc analysis. p < 0.05 was considered statistically significant. P ≥ 0.055 to p ≤ 0.104 was considered close-to-significant (CS).

## ACKNOWLEDGMENTS

We thank Nancy Busija, MA, for editing the manuscript. We thank Dana Liu for technical help.

## AUTHORS’ CONTRIBUTIONS

S.C., P.K.C, and D.W.B. conceived and designed the experiments; S.C., P.K.C., J.C.H., I.R., and J.J.G. performed experiments; S.C., P.K.C., J.C.H., and J.J.G analyzed data, interpreted experimental results, and prepared figures; S.C., P.K.C., and J.C.H. drafted the manuscript; S.C., P.K.C., J.C.H., I.R., J.J.G., P.V.G.K., J.M.G., and D.W.B. edited and revised the manuscript and approved the final version of the manuscript.

## Summary Figure

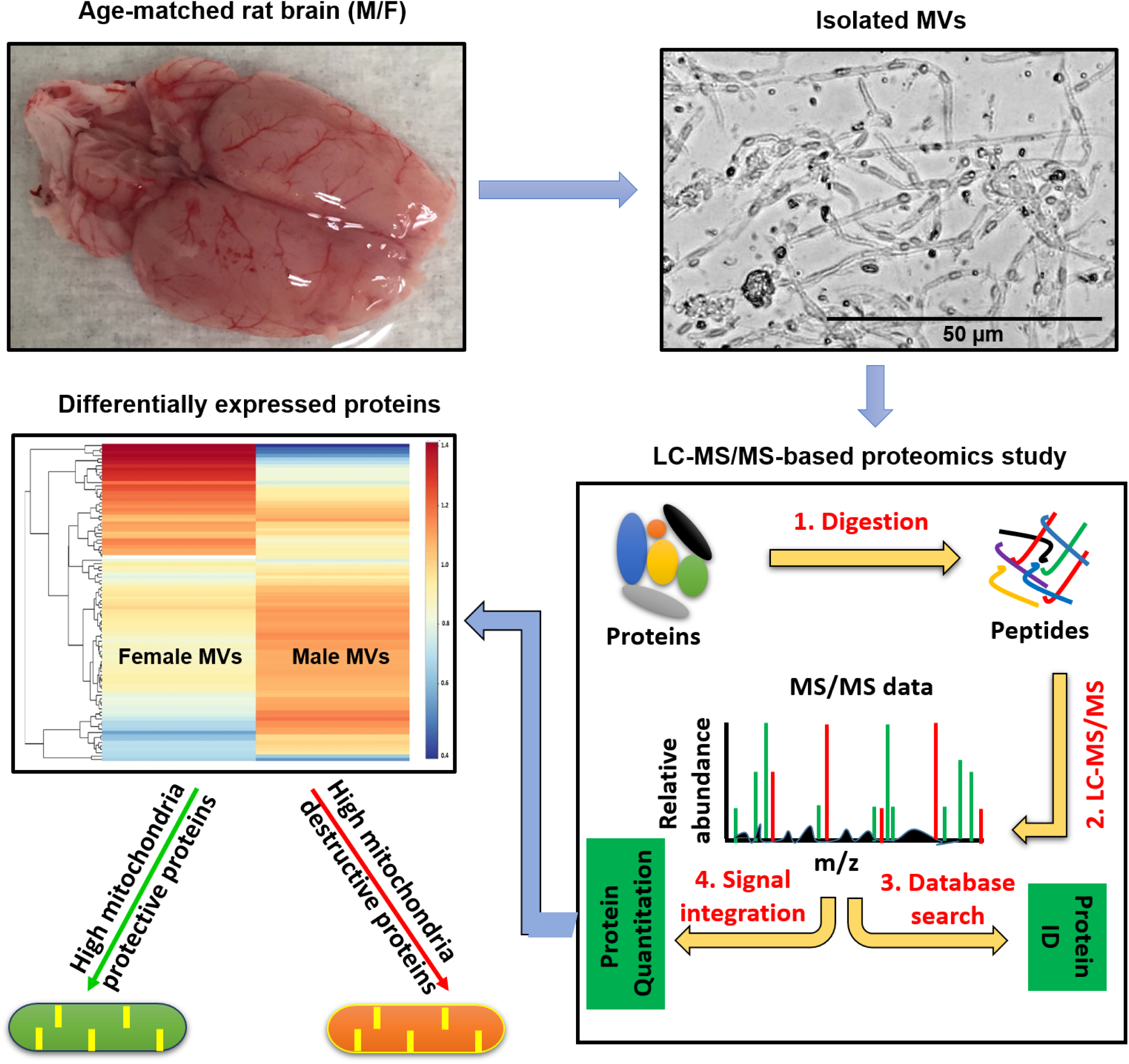

Mitochondrial characteristics in the male and female cerebral microcirculation were examined since evidence suggests that metabolic status of this vascular segment is a major determinant of neurological pathologies such as cognitive impairment, vascular dementia, and AD and because the occurrence and progression of these neurological diseases vary between the sexes. Major findings are that:

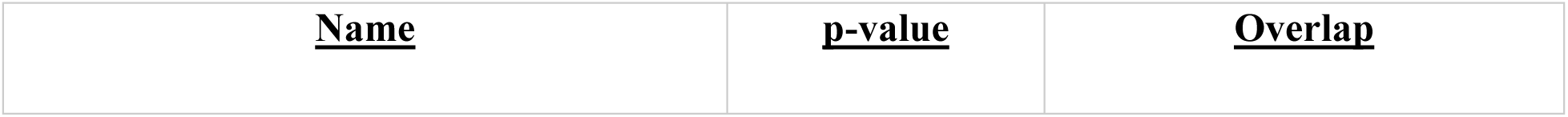

- Female MVs have a greater abundance of mitochondrial structural and energy producing proteins as well as more anti-inflammatory/pro-healing proteins associated with mitochondrial activity than males.
- Male MVs have a higher abundance of mitochondrial destructive proteins than females.
- Female MVs exhibit greater a potential for using alternative fuels than males.

## REFERENCES

Bachar AR, Scheffer L, Schroeder AS, Nakamura HK, Cobb LJ, Oh YK, Lerman LO, Pagano RE, Cohen P, Lerman A (2010) Humanin is expressed in human vascular walls and has cytoprotective effect against oxidized LDL-induced oxidative stress. Cardiovasc Res 88:360–366.

Banerjee P, Chander V, Bandyopadhyay A (2015). Balancing functions of annexin A6 maintain equilibrium between hypertrophy and apoptosis in cardiomyocytes. Cell Death Dis 6: e1873.

Bourassa P, Tremblay C, Schneider JA, Bennett DA, Calon F (2019). Beta-amyloid pathology in human brain microvessel extracts from the parietal cortex: relation with cerebral amyloid angiopathy and Alzheimer’s disease. Acta Neuropathol 137: 801–823.

Brzica H, Abdullahi W, Reilly BG, Ronaldson PT (2018). Sex-specific differences in organic anion transporting polypeptide 1a4 (Oatp1a4) functional expression at the blood-brain barrier in Sprague-Dawley rats. Fluids Barriers CNS 15:25.

Burstein SR, Kim HJ, Fels JA, Qian L, Zhang S, Zhou P, Starkov AA, Iadecola C, Manfredi G (2018). Estrogen receptor beta modulates permeability transition in brain mitochondria. Biochim Biophys Acta Bioenerg 1859: 423–433.

Busija DW and Katakam PV (2014) Mitochondrial mechanisms in cerebral vascular control: shared signaling pathways with preconditioning. J Vasc Res 51: 175–189.

Busija DW, Rutkai I, Dutta S, Katakam PV (2016) Role of mitochondria in cerebral vascular function: energy production, cellular protection, and regulation of vascular tone. Compr Physiol 6: 1529–1548.

Capone C, Anrather J, Milner TA, Iadecola C (2009). Estrous cycle-dependent neurovascular dysfunction induced by angiotensin II in the mouse neocortex. Hypertension 54: 302–307.

Castaldo P, Cataldi M, Magi S, Lariccia V, Arcangeli S, Amoroso S (2009). Role of the mitochondrial sodium/calcium exchanger in neuronal physiology and in the pathogenesis of neurological diseases. Prog Neurobiol 87: 58–79.

Chauhan A, Moser H, McCullough LD (2017). Sex differences in ischaemic stroke: potential cellular mechanisms. Clin Sci (Lond) 131: 533–552.

Chlystun M, Campanella M, Law AL, Duchen MR, Fatimathas L, Levine TP, Gerke V, Moss SE (2013). Regulation of mitochondrial morphogenesis by annexin A6. PLoS One 8: e53774.

Dai DF, Rabinovitch PS, Ungvari Z (2012) Mitochondria and cardiovascular aging. Circ Res 110:1109– 1124.

de Souza Mota C, Weis SN, Almeida RF, Dalmaz C, Guma FTC, Pettenuzzo LF (2017). Chronic Stress Causes Sex-Specific and Structure-Specific Alterations in Mitochondrial Respiratory Chain Activity in Rat Brain. Neurochem Res 42: 3331–3340.

Demarest TG, Schuh RA, Waddell J, McKenna MC, Fiskum G (2016). Sex-dependent mitochondrial respiratory impairment and oxidative stress in a rat model of neonatal hypoxic-ischemic encephalopathy. J Neurochem 137: 714–29.

Demarest TG, Schuh RA, Waite EL, Waddell J, McKenna MC, Fiskum G (2016). Sex dependent alterations in mitochondrial electron transport chain proteins following neonatal rat cerebral hypoxic-ischemia. J Bioenerg Biomembr 48: 591–598.

Díaz A, López-Grueso R, Gambini J, Monleón D, Mas-Bargues C, Abdelaziz KM, Viña J, Borrás C (2019). Sex Differences in Age-Associated Type 2 Diabetes in Rats-Role of Estrogens and Oxidative Stress. Oxid Med Cell Longev 2019:6734836.

Eliceiri BP, Gonzalez AM, Baird A (2011). Zebrafish model of the blood-brain barrier: morphological and permeability studies. Methods Mol Biol 686: 371–378.

Erdő F, Krajcsi P (2019). Age-Related Functional and Expressional Changes in Efflux Pathways at the Blood-Brain Barrier. Front Aging Neurosci 11: 196.

Giatromanolaki A, Sivridis E, Mendrinos S, Koutsopoulos AV, Koukourakis MI (2014). Autophagy proteins in prostate cancer: relation with anaerobic metabolism and Gleason score. Urol Oncol 32: 39.e11–8.

Gomez-Zepeda D, Taghi M, Smirnova M, Sergent P, Liu WQ, Chhuon C, Vidal M, Picard M, Thioulouse E, Broutin I, Guerrera IC, Scherrmann JM, Parmentier Y, Decleves X, Menet MC (2019). LC-MS/MS-based quantification of efflux transporter proteins at the BBB. J Pharm Biomed Anal 164: 496–508.

Grimmig B, Kim SH, Nash K, Bickford PC, Douglas SR (2017) Neuroprotective mechanisms of astaxanthin: a potential therapeutic role in preserving cognitive function in age and neurodegeneration. Geroscience 39: 19–32.

Hansra GK, Popov G, Banaczek PO, Vogiatzis M, Jegathees T, Goldbury CS, Cullen KM (2019). The neuritic plaque in Alzheimer’s disease: perivascular degeneration of neuronal processes. Neurobiol Aging 82: 88–101.

Harman JC, Guidry JJ, Gidday JM (2018) Comprehensive characterization of the adult ND4 Swiss Webster mouse retina: Using discovery-based mass spectrometry to decipher the total proteome and phosphoproteome. Mol Vis 24: 875–889.

Huang X, Bai HM, Chen L, Li B, Lu YC (2010). Reduced expression of LC3B-II and Beclin 1 in glioblastoma multiforme indicates a down-regulated autophagic capacity that relates to the progression of astrocytic tumors. J Clin Neurosci 17: 1515–1519.

Iadecola C, Duering M, Hachinski V, Joutel A, Pendlebury ST, Schneider JA, Dichgans M (2019). Vascular Cognitive Impairment and Dementia: JACC Scientific Expert Panel. J Am Coll Cardiol 73: 3326–3344.

Javitt A, Merbl Y (2019). Global views of proteasome-mediated degradation by mass spectrometry. Expert Rev Proteomics 16:711–716.

Johnson AC, Cipolla MJ (2015). The cerebral circulation during pregnancy: adapting to preserve normalcy. Physiology (Bethesda) 30: 139–147.

Kabeya Y, Mizushima N, Ueno T, Yamamoto A, Kirisako T, Noda T, Kominami E, Ohsumi Y, Yoshimori T (2000). LC3, a mammalian homologue of yeast Apg8p, is localized in autophagosome membranes after processing. EMBO J 19: 5720–5728.

Kemper MF, Zhao Y, Duckles SP, Krause DN (2013) Endogenous ovarian hormones affect mitochondrial efficiency in cerebral endothelium via distinct regulation of PGC-1 isoforms. J Cereb Blood Flow Metab 33: 122–128.

Kemper MF, Stirone C, Krause DN, Duckles SP, Procaccio V (2014) Genomic and non-genomic regulation of PGC1 isoforms by estrogen to increase cerebral vascular mitochondrial biogenesis and ROS protection. Eur J Pharmacol 723: 322–329.

Kitamura N, Nakamura Y, Miyamoto Y, Miyamoto T, Kabu K, Yoshida M, Futamura M, Ichinose S, Arakawa H (2011). Mieap, a p53-inducible protein, controls mitochondrial quality by repairing or eliminating unhealthy mitochondria. PLoS One 6: e16060.

Ko YH, Cho YS, Won HS, Jeon EK, An HJ, Hong SU, Park JH, Lee MA (2013). Prognostic significance of autophagy-related protein expression in resected pancreatic ductal adenocarcinoma. Pancreas 42: 829–835.

Kowalczyk JE, Beresewicz M, Gajkowska B, Zabłocka B (2009). Association of protein kinase C delta and phospholipid scramblase 3 in hippocampal mitochondria correlates with neuronal vulnerability to brain ischemia. Neurochem Int 55: 157–63.

Lazova R, Camp RL, Klump V, Siddiqui SF, Amaravadi RK, Pawelek JM (2012). Punctate LC3B expression is a common feature of solid tumors and associated with proliferation, metastasis, and poor outcome. Clin Cancer Res 18: 370–379.

Lee ES, Yoon JH, Choi J, Andika FR, Lee T, Jeong Y (2019a). A mouse model of subcortical vascular dementia reflecting degeneration of cerebral white matter and microcirculation. J Cereb Blood Flow Metab 39: 44–57.

Lee J, Pinares-Garcia P, Loke H, Ham S, Vilain E, Harley VR (2019b). Sex-specific neuroprotection by inhibition of the Y-chromosome gene, SRY, in experimental Parkinson’s disease. Proc Natl Acad Sci U S A 116: 16577–16582.

Liu J, Dai Q, Chen J, Durrant D, Freeman A, Liu T, Grossman D, Lee RM (2003). Phospholipid scramblase 3 controls mitochondrial structure, function, and apoptotic response. Mol Cancer Res 1: 892–902.

Liu Y, Braidy N, Poljak A, Chan DKY, Sachdev P (2018). Cerebral small vessel disease and the risk of Alzheimer’s disease: A systematic review. Ageing Res Rev 47: 41–48.

Merdzo I, Rutkai I, Sure VN, McNulty CA, Katakam PV, Busija DW (2017). Impaired Mitochondrial Respiration in Large Cerebral Arteries of Rats with Type 2 Diabetes. J Vasc Res 54:1–12.

Merdzo I, Rutkai I, Tokes T, Sure VN, Katakam PV, Busija DW (2016). The mitochondrial function of the cerebral vasculature in insulin-resistant Zucker obese rats. Am J Physiol Heart Circ Physiol 310: H830–838.

Mizushima N, Noda T, Yoshimori T, Tanaka Y, Ishii T, George MD, Klionsky DJ, Ohsumi M, Ohsumi Y (1998). A protein conjugation system essential for autophagy. Nature 395: 395–398.

Mizushima N, Yamamoto A, Hatano M, Kobayashi Y, Kabeya Y, Suzuki K, Tokuhisa T, Ohsumi Y, Yoshimori T (2001). Dissection of autophagosome formation using Apg5-deficient mouse embryonic stem cells. J Cell Biol 152: 657–668.

Nakamura Y, Arakawa H (2017). Discovery of Mieap-regulated mitochondrial quality control as a new function of tumor suppressor p53. Cancer Sci 108: 809–817.

Netto CA, Sanches E, Odorcyk FK, Duran-Carabali LE, Weis SN (2017). Sex-dependent consequences of neonatal brain hypoxia-ischemia in the rat. J Neurosci Res 95: 409–421.

Porte B, Chatelain C, Hardouin J, Derambure C, Zerdoumi Y, Hauchecorne M, Dupré N, Bekri S, Gonzalez B, Marret S, Cosette P, Leroux P (2017). Proteomic and transcriptomic study of brain microvessels in neonatal and adult mice. PLoS One 12: e0171048.

Reddy PH, Beal MF (2008) Amyloid beta, mitochondrial dysfunction and synaptic damage: implications for cognitive decline in aging and Alzheimer’s disease. Trends Mol Med 14:45–53.

Romano A, Koczwara JB, Gallelli CA, Vergara D, Micioni Di Bonaventura MV, Gaetani S, Giudetti AM (2017). Fats for thoughts: An update on brain fatty acid metabolism. Int J Biochem Cell Biol 84: 40–45.

Rutkai I, Dutta S, Katakam PV, Busija DW (2015). Dynamics of enhanced mitochondrial respiration in female compared with male rat cerebral arteries. Am J Physiol Heart Circ Physiol 309: H1490–1500.

Rutkai I, Merdzo I, Wunnava S, Curtin G, Katakam PVG, Busija DW (2017) Cerebrovascular function and mitochondrial bioenergetics after ischemia-reperfusion in male rats. J Cereb Blood Flow Metab 39:1056–1068.

Silbergeld DL, Ali-Osman F (1991). Isolation and characterization of microvessels from normal brain and brain tumors. J Neurooncol 11(1): 49–55.

Snowden SG, Ebshiana AA, Hye A, An Y, Pletnikova O, O’Brien R, Troncoso J, Legido-Quigley C, Thambisetty M (2017). Association between fatty acid metabolism in the brain and Alzheimer disease neuropathology and cognitive performance: A nontargeted metabolomic study. PLoS Med 14: e1002266.

Sure VN, Sakamuri SSVP, Sperling JA, Evans WR, Merdzo I, Mostany R, Murfee WL, Busija DW, Katakam PVG (2018) A novel high-throughput assay for respiration in isolated brain microvessels reveals impaired mitochondrial function in the aged mice. Geroscience 40: 365–375.

Tang BL (2015). MIRO GTPases in Mitochondrial Transport, Homeostasis and Pathology. Cells 5: 1.

Tharakan R, Kreimer S, Ubaida-Mohien C, Lavoie J, Olexiouk V, Menschaert G, Ingolia NT, Cole RN, Ishizuka K, Sawa A, Nucifora LG (2019). A methodology for discovering novel brain-relevant peptides: Combination of ribosome profiling and peptidomics. Neurosci Res doi: 10.1016/j.neures.2019.02.006.

Ungvari Z, Sonntag WE, Csiszar A (2010) Mitochondria and aging in the vascular system. J Mol Med (Berl) 88:1021–1027.

van Leijsen EMC, de Leeuw FE, Tuladhar AM (2017). Disease progression and regression in sporadic small vessel disease-insights from neuroimaging. Clin Sci (Lond) 131: 1191–1206.

Vandenbrouck Y, Christiany D, Combes F, Loux V, Brun V (2019). Bioinformatics Tools and Workflow to Select Blood Biomarkers for Early Cancer Diagnosis: An Application to Pancreatic Cancer. Proteomics e1800489.

Wardlaw JM, Smith C, Dichgans M (2019). Small vessel disease: mechanisms and clinical implications. Lancet Neurol 18: 684–696.

Werle K, Chen J, Xu HG, Zhao RX, He Q, Lu C, Cui R, Liang J, Li YL, Xu ZX (2014). Liver kinase B1 regulates the centrosome via PLK1. Cell Death Dis 5: e1157.

Yen K, Lee C, Mehta H, Cohen P (2013) The emerging role of the mitochondrial-derived peptide humanin in stress resistance. J Mol Endocrin 50: R11–R19.

Yousif S, Marie-Claire C, Roux F, Scherrmann JM, Declèves X (2007). Expression of drug transporters at the blood-brain barrier using an optimized isolated rat brain microvessel strategy. Brain Res 1134: 1–11.

Yu H, Yang C, Chen S, Huang Y, Liu C, Liu J, Yin W (2017) Comparison of the glycopattern alterations of mitochondrial proteins in cerebral cortex between rat Alzheimer’s disease and the cerebral ischemia model. Sci Rep 7:39948.

Zhang F, Wang W, Siedlak SL, Liu Y, Liu J, Jiang K, Perry G, Zhu X, Wang X (2015). Miro1 deficiency in amyotrophic lateral sclerosis. Front Aging Neurosci 7:100.

